# Genomic diversity and evolution of coronavirus (SARS-CoV-2) in France from 309 COVID-19-infected patients

**DOI:** 10.1101/2020.09.04.282616

**Authors:** Anthony Levasseur, Jeremy Delerce, Aurelia Caputo, Ludivine Brechard, Philippe Colson, Jean-Christophe Lagier, Pierre-Edouard Fournier, Didier Raoult

## Abstract

The novel coronavirus (SARS-CoV-2) causes pandemic of viral pneumonia. The evolution and mutational events of the SARS-CoV-2 genomes are critical for controlling virulence, transmissibility, infectivity, severity of symptoms and mortality associated to this infectious disease. We collected and investigated 309 SARS-CoV-2 genomes from patients infected in France. Detailed genome cartography of all mutational events (SNPs, indels) was reported and correlated to clinical features of patients. A comparative analysis between our 309 SARS-CoV-2 genomes from French patients and the reference Wuhan coronavirus genome revealed 315 substitution mutations and six deletion events: ten were in 5’/3’ UTR, 178 were nonsynonymous, 126 were synonymous and one generated a stop codon. Six different deleted areas were also identified in nine viral variants. In particular, 30 substitution mutations (18 nonsynonymous) and one deletion (Δ21765-21770) concerned the spike S glycoprotein. An average of 7.8 mutational events (+/- 1.7 SD) and a median of 8 (range, 7-9) were reported per viral isolate. Comparative analyses and clustering of specific mutational signatures in 309 genomes disclose several divisions in groups and subgroups combining their geographical and phylogenetic origin. Clinical outcomes of the 309 COVID-19-infected patients were investigated according to the mutational signatures of viral variants. These findings highlight the genome dynamics of the coronavirus 2019-20 and shed light on the mutational landscape and evolution of this virus. Inclusion of the French cohort enabled us to identify 161 novel mutations never reported in SARS-CoV-2 genomes collected worldwide. These results support a global and continuing surveillance of the emerging variants of the coronavirus SARS-CoV-2.

## INTRODUCTION

The new coronavirus SARS-CoV-2 has now spread to every country in the world. SARS-CoV-2 is classified among the positive-sense single-stranded (ss)RNA viruses as the seventh coronavirus known to infect human^1^. Four genera of coronaviruses have been identified in the subfamily Orthocoronavirinae of the Coronaviridae family (order Nidovirales). Described for the first time in December 2019 in Wuhan, China, the new SARS-CoV-2 coronavirus will naturally accumulate and fix mutations in its RNA genome as the epidemic evolves, partly due to the high immunological pressure in humans. A number of characteristics of coronaviruses are still unknown and one of the key questions is how fast is the SARS-CoV-2 virus evolving? Indeed, mutations could drastically result in difference in virulence, transmissibility, and clinical evolution. Therefore, the frequency and pattern of mutations are essential features to be considered in this ongoing global epidemic.

The genome of SARS-CoV-2 is a positive ssRNA of approximately 29.9 kbases in length, with a 5’cap structure and 3’polyadenine tail^2^. Two large ORFs are translated into polyproteins arising from ribosomal frameshift that are subsequently post-translationally processed to produce 16 proteins conserved between coronaviruses^3^. These 16 proteins are involved in the synthesis of viral RNA and immune evasion^4^. Four structural proteins (spike S, envelope E, membrane M and nucleocapsid N) and at least nine small accessory proteins, including proteins unique to SARS-CoV-2, are produced during viral genome replication^3,5^.

By comparing alpha- and beta-coronaviruses, it appears that SARS-CoV-2 exhibits notable genomic features such as an optimized binding to the human receptor ACE2 and a functional polybasic (furin) site in the spike protein with predicted O-linked glycans^6^. The receptor binding domain (RBD) exhibits the most variable fraction of the genome of SARS-CoV-2 (only ∼40% amino acid identity with other SARS-CoV coronaviruses). The origin of this virus is debated but two main hypotheses emerged based on either the natural selection in an animal host before zoonotic transfer or natural selection in humans following zoonotic transfer^7^. Nearly identical RBDs found in pangolins coronaviruses and SARS-CoV-2 could indicate an acquisition in SARS-CoV-2 by recombination and mutations in a parsimonious scenario^6^. Metagenomic sequencing identified coronaviruses close to SARS-CoV-2 from Malayan pangolin and confirmed the natural reservoir of this virus in wild animals and the risk of zoonotic transmission^8^.

Being a RNA virus, SARS-CoV-2 is prone to mutate because of the lack of proofreading activity of polymerase. Previous work in SARS-CoV reported an exoribonuclease domain (ExoN) in non-structural protein 14 that could provide proofreading activity protecting the virus from high rate of mutagenesis^9^. Mutations have to be carefully studied and related to the severity of symptoms and clinical evolution associated to patients. In addition, the mutation rate correlates with drug resistance and escape from immune surveillance. The emerging respiratory disease outbreaks caused by the SARS-CoV-2 required an urgent need for comprehensive studies that combine genomic data, epidemiological data, and chart records of the clinical symptoms and clinical evolution of patients. In addition to genomic mutations, transcriptomic studies have revealed several viable deletions in essential proteins, including glycoprotein S reinforcing that several regions are prone to mutate^10^.

The 2019–20 coronavirus pandemic was confirmed to have spread to France since January 2020. In this work, 309 genomes of SARS-CoV-2 from French patients monitored in IHU Méditerranée Infection were studied and compared to the isolates from different countries in all continents. Mutational events were evidenced and compared to the reference Wuhan coronavirus isolate. Finally, clinical issues and impacts of these mutations were investigated.

## RESULTS

### Mutational landscape of the SARS-CoV-2 genomes from the French cohort

The mutational landscape of 309 coronavirus genomes was tracked in terms of SNPs and insertions/deletions (Table 1). As compared to the reference genome SARS-CoV-2 isolated in Wuhan in December 2019, a total of 321 mutational events were reported in the SARS-CoV-2 genomes from 309 patients. A mean of 7.8 mutational nucleotide substitutions (+/- 1.7 SD) and a median of 8 (range, 7-9) were reported per viral isolate. Ten substitutions were retrieved in the non-coding part of the genome. In the coding region, 305 nucleotide substitutions were detected representing 178 nonsynonymous (∼58.4%) and 126 synonymous mutations (∼41.3%) and one stop codon mutation (∼0.3%). Some mutations were commonly distributed in the majority of our isolates whereas 208 substitutions (∼68.2%) were unique to one isolate. Regarding missense mutations, we noted drastic changes of aminoacid (AA) in which the physicochemical properties of the AA were severely interfered (Table 2). In particular, the spike gene contained 30 substitutions (18 nonsynonymous, 60%) and included one deletion of six nucleotides (Δ21765-21770). Among mutations retrieved in the spike gene, the missense mutation P330S (22550, CCT>TCT) was embedded at the border of the receptor-binding domain (RBD). A stop mutation was also found in ORF7a at the protein position 38, meaning that the protein was prematurely truncated at the first third of its sequence.

**Table 1.**
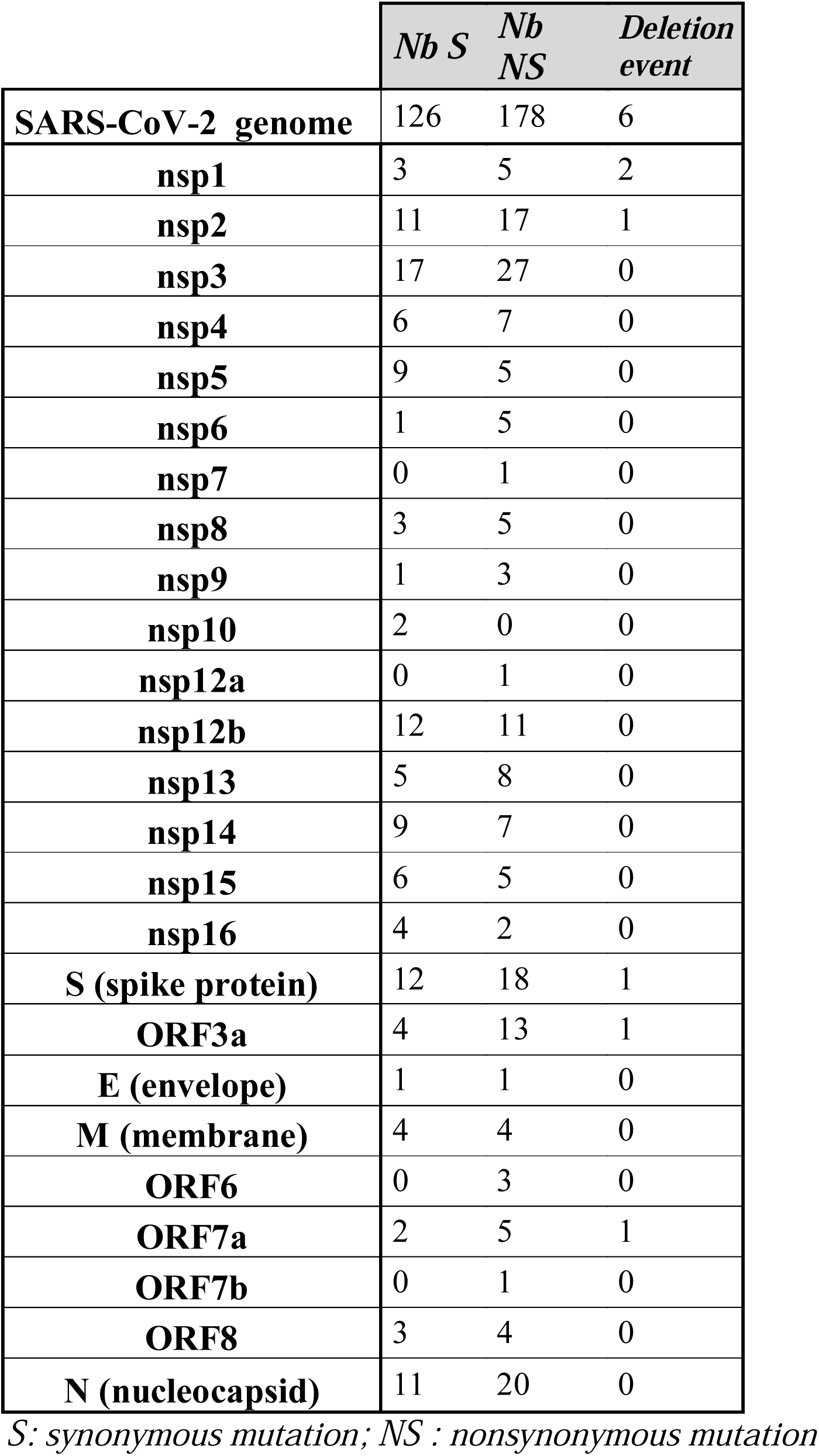
Mutational events detected in the 309 SARS-CoV-2 genomes as compared to the reference Wuhan coronavirus.

**Table 2.**
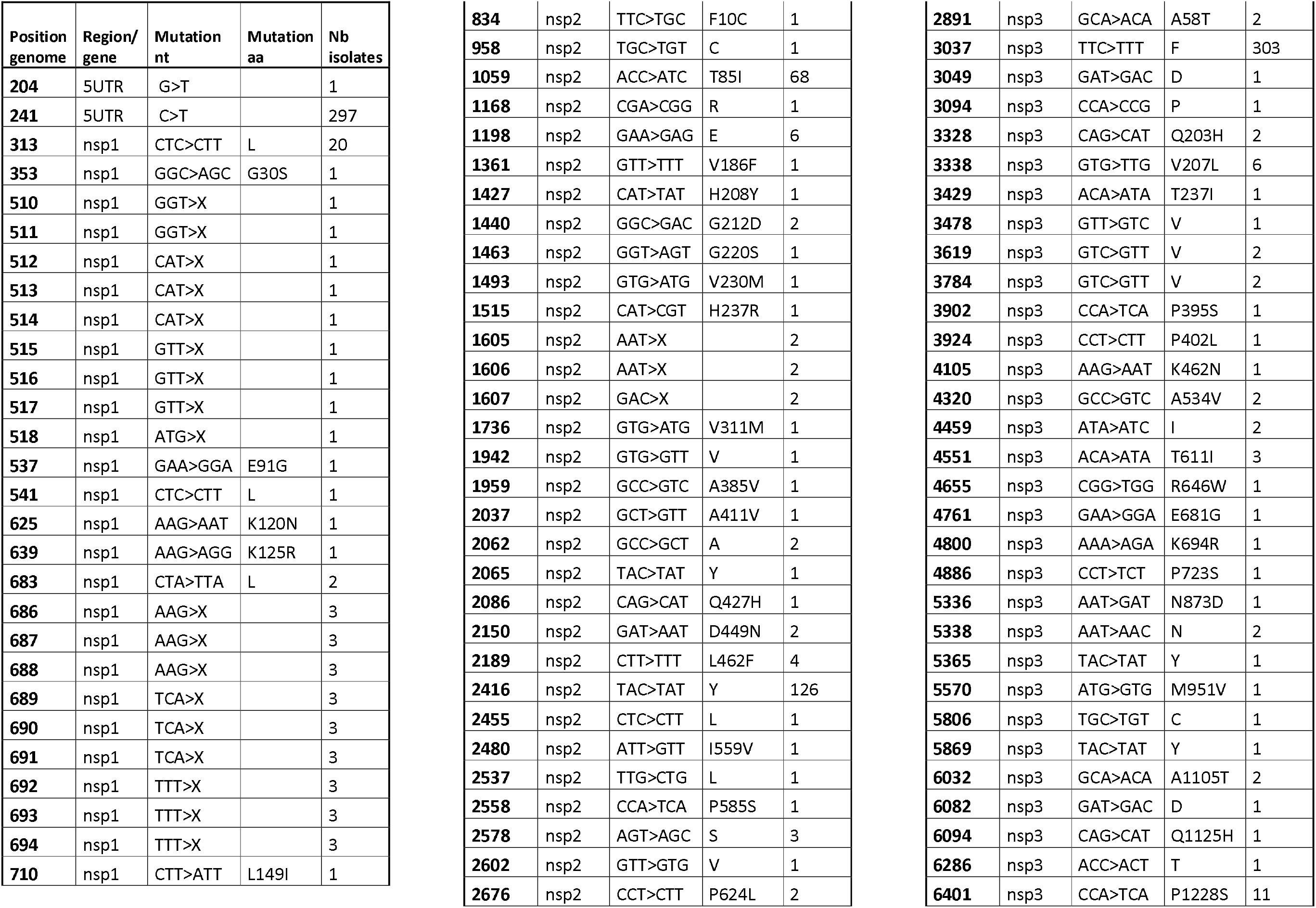
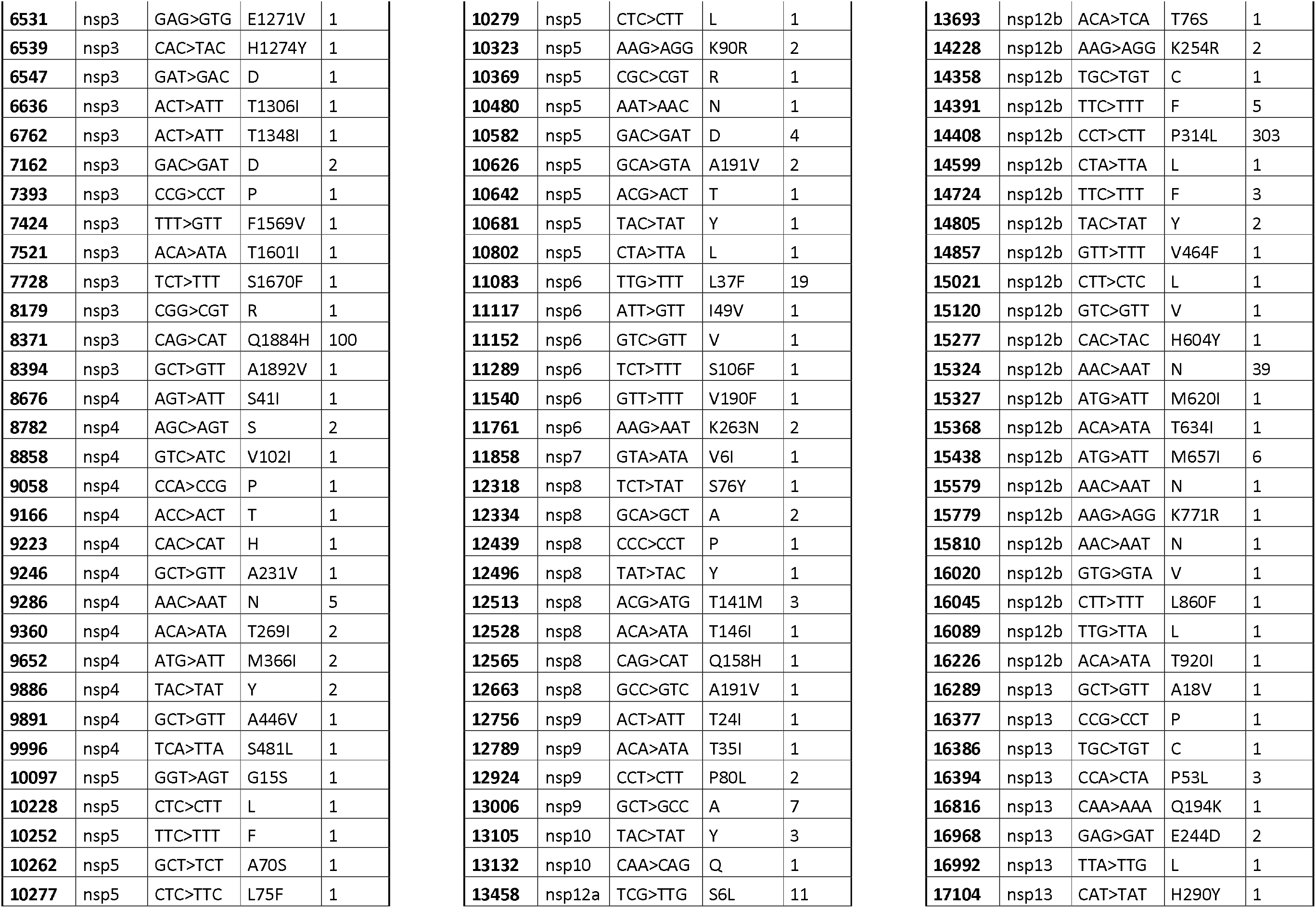
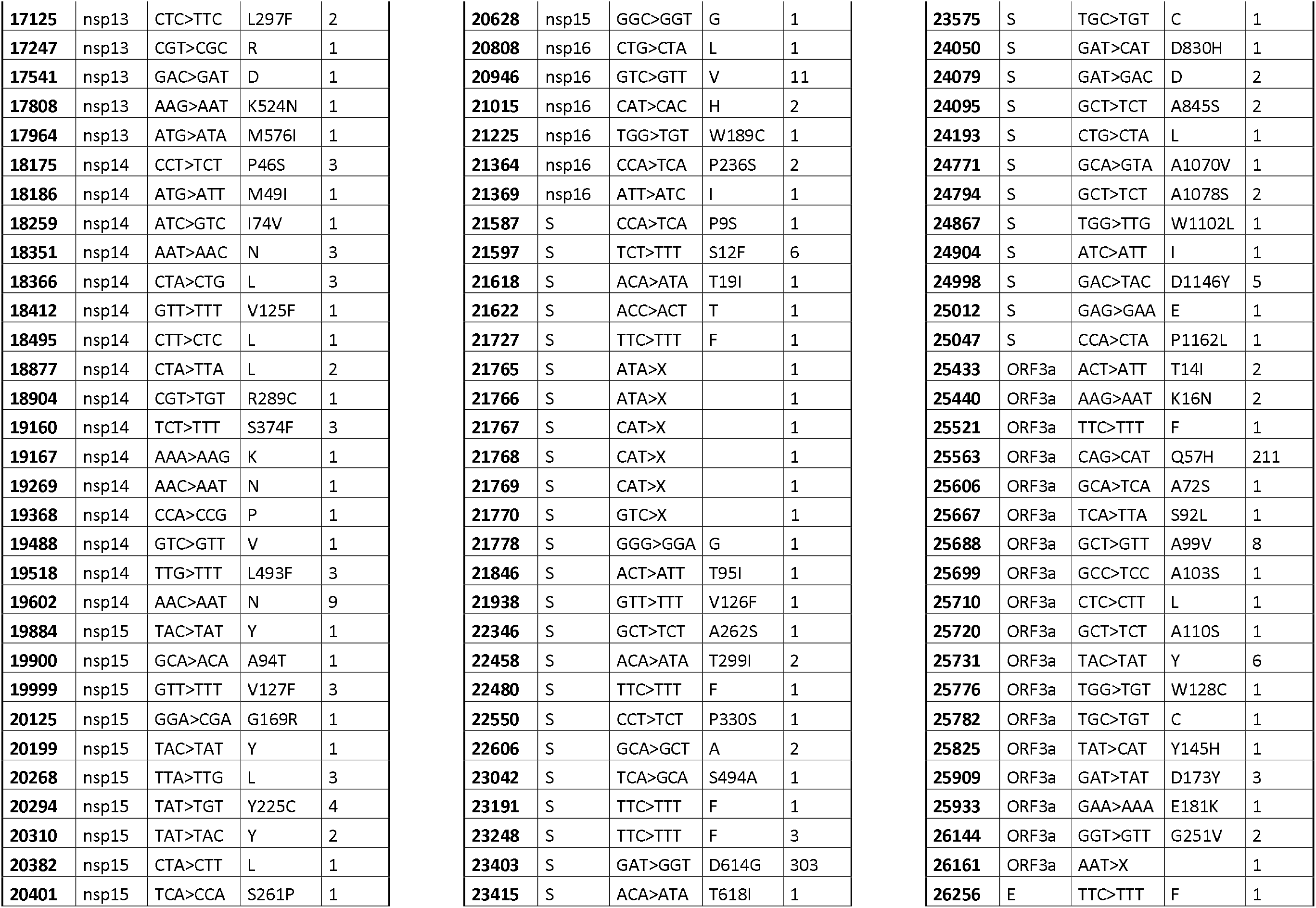
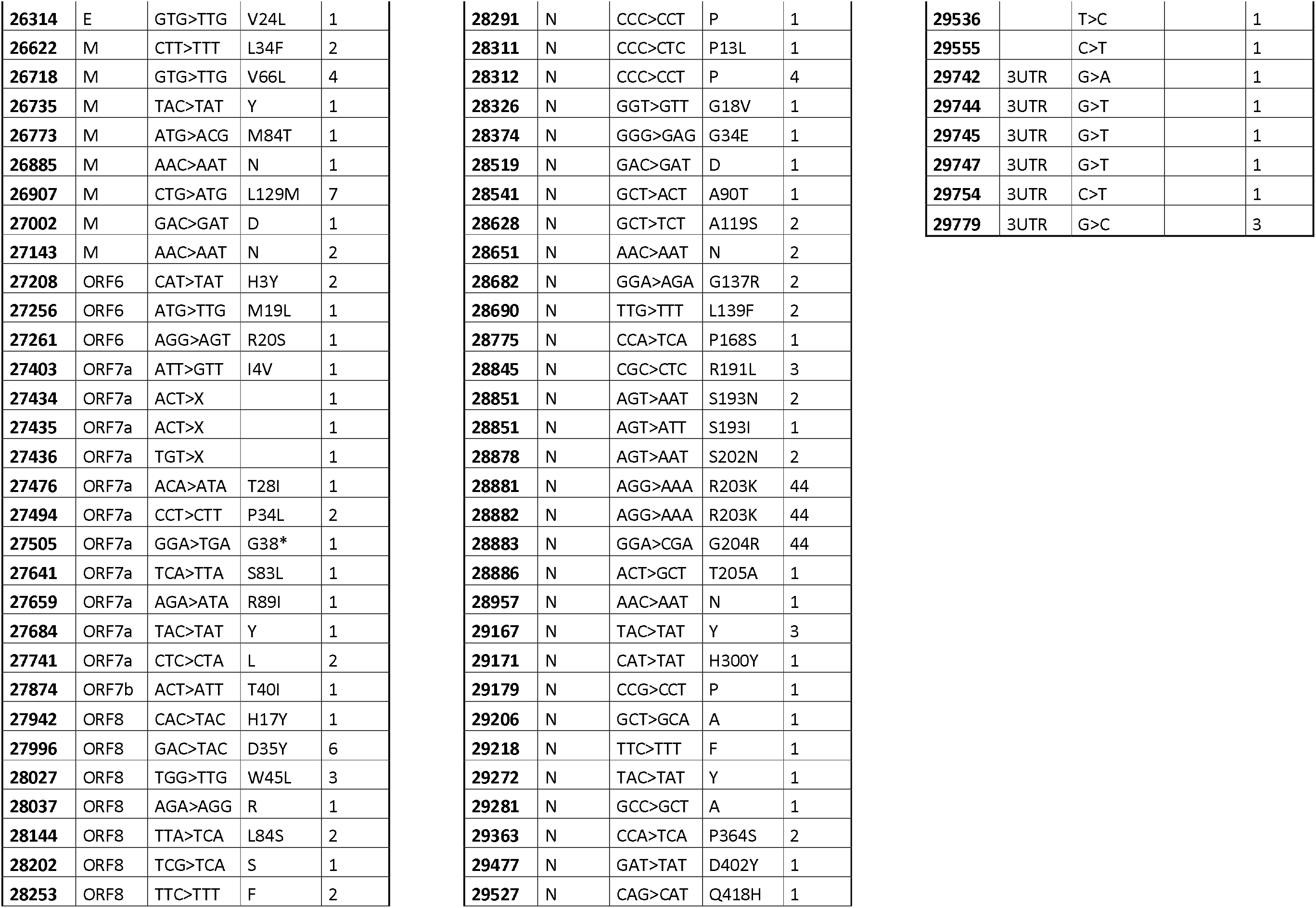
Detailed mutations detected in the 309 SARS-CoV-2 genomes from French patients

Interestingly, we identified six putative deleted regions distributed in nine viral variants among the genes nsp1, nsp2, spike S, ORF3a and ORF7a. These deletions removed entire codons for nsp1 (Δ510-518 and Δ686-694, three AA), nsp2 (Δ1605-1607, one AA), spike protein (Δ21765-21770, two AA), ORF7a (Δ27434-27436, one AA) and consequently, conserved open reading frames. Moreover, the deletion event retrieved in nsp1 (position Δ686-694) was found in three different patients and the deletion in nsp2 (position Δ1605-1607) in two different patients that were not epidemiologically-related, meaning that these two evolutionary events could be a convergent evolution of the virus. In addition, one single nucleotide deletion impacted ORF3a at the position Δ26161, this frameshift mutation lead to a truncated protein missing 19 AA with a stop codon generated at the protein position 259.

The occurrence of specific mutations in the 309 genomes was studied and classified. In order to organize our dataset into meaningful groups and identify coherent patterns, a double hierarchical clustering of the detected mutations was generated (Figure 1). The hierarchical clustering revealed preponderant mutations shared by a majority of the isolates. It divided our 309 patients into five main clusters corresponding to specific mutational signatures. Regarding the clustering of mutational events, we detected the presence and co-occurrence of specific mutations in different clusters (Figure 1). Among the main clusters, we pointed the cluster 1 (44 patients, 14.2%, positions [28881-28882-28883]) with two nonsynonymous mutations in protein N (R203K; G204R). Cluster 2 (39 patients, 12.6%, position 15324) contains a synonymous mutation (C15324T). Cluster 3 (126, 100 and 211 patients, at positions 2416, 8371, 25563, respectively) includes one synonymous mutation (C2416T), and two nonsynonymous mutations (nsp3: Q1884H; ORF3a: Q57H). Cluster 4 (68 patients, 22%, position 1059) contains one nonsynonymous mutation (nsp2: T85I). Finally, cluster 5 (from 297 to 303 patients, 96-98%, positions 241, 3037, 14408, 23403) displays one mutation in 5’UTR (C241T), one synonymous mutation (C3037T) and two nonsynonymous mutations (nsp12b: P314L, S protein D614G).

**Figure 1.**
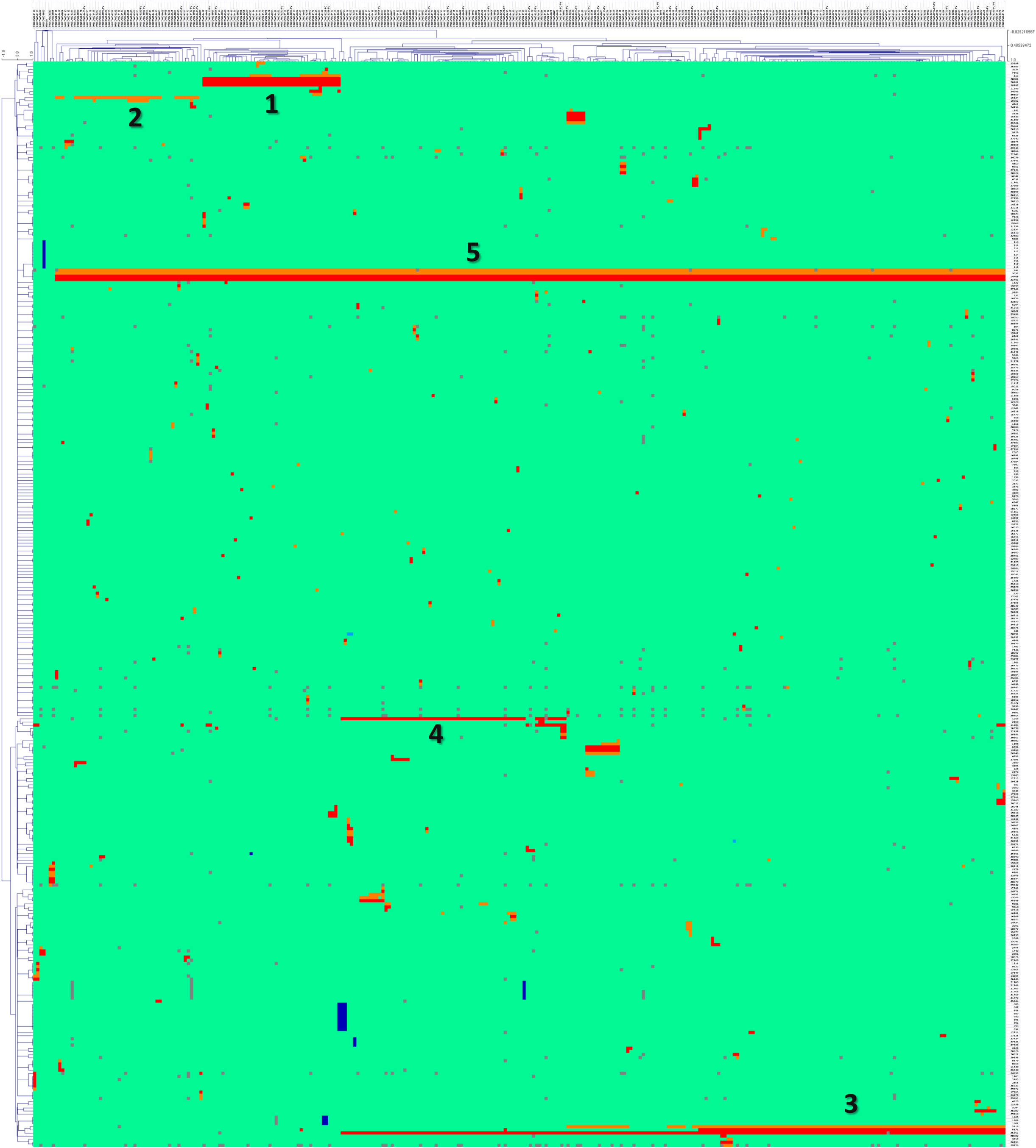
Double hierarchical clustering of mutational events in SARS-CoV-2 genomes collected in 309 COVID-19 infected French patients. Top tree: Viral isolates from 309 patients. Left tree: Position of identified mutations according to the reference Wuhan genome. Clusters (1 to 5) were labeled with numbers ranging from 1 to 5. Missing positions were labeled in grey. The mutation type is represented by a colour scale from green (position identical to the reference), red (nonsynonymous mutation), orange (synonymous mutation) and blue (deleted position).

The genomes of all 309 isolates were integrated in a global phylogenetic reconstruction including all other 200 genomes of SARS-CoV-2 available in France (Figure 2 + supplementary figure 1). We observed different clades corresponding to the previous mutational patterns revealed by the hierarchical clustering. These mutations were positioned in phylogenetic trees. Globally, the distribution of the 309 SARS-CoV-2 genomes from our patients was distributed across all lineages. Noteworthy, we observed a high density of isolates from our cohort in the cluster 3a corresponding to the mutation nsp3: G8371T, Q1884H. Phylogenetic reconstruction was also carried out including all complete genomes available worldwide (5519 genomes on 15^th^ April 2020). All viral isolates of our French cohort were mainly distributed in clades including European isolates (Figure 3). Again, the mutation in the cluster 3a (nsp3: G8371T, Q1884H) regrouped almost exclusively the variants from our French cohort (98/105 genomes). Compared to the Chinese reference, we observed a prevalent distribution of specific mutations at positions 241, 3037, 14808 and 23403 that spread in approximately ∼3570 (+/-7) genomes representing ∼64.6 % of the isolates worldwide. These mutations have been reported in variants from all continents. Although, we detected some clades depending on the common geographical origin of the patients, a global distribution of variants in all continents was observed, therefore meaning and confirming that SARS-CoV-2 is circulating worldwide.

**Figure 2.**
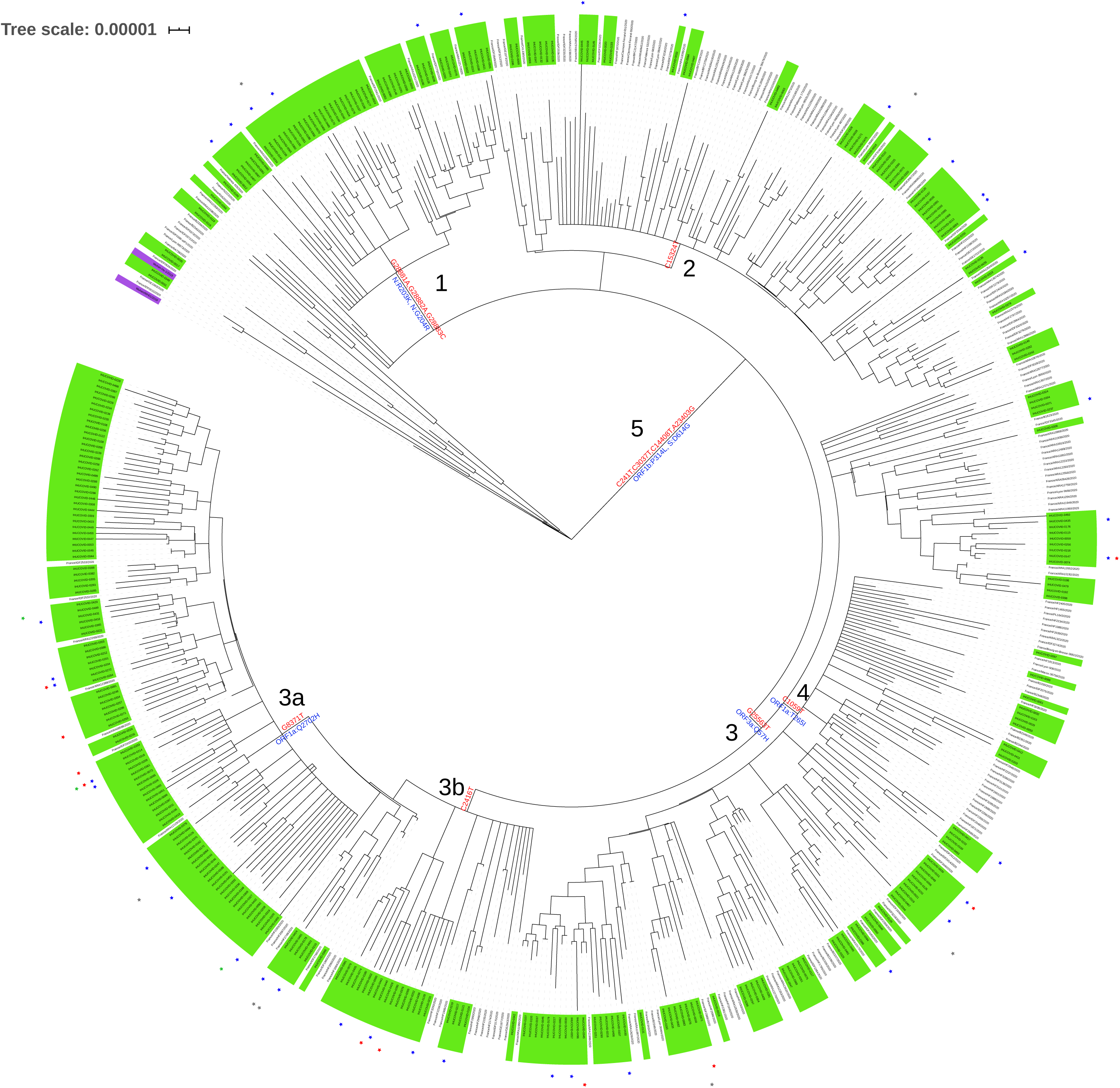
Phylogenetic tree of coronavirus in France. The phylogenetic tree was reconstructed from 509 viral genomes collected in France (309 from this study). Number represents the cluster identified in the double hierarchical clustering. The reference Wuhan coronavirus isolate was underlined in purple. Red star corresponds to PClinO, blue star corresponds to PVirO, grey star corresponds to the loss of follow-up and green star corresponds to deceased patient. Main mutations were labeled along branches.

**Figure 3.**
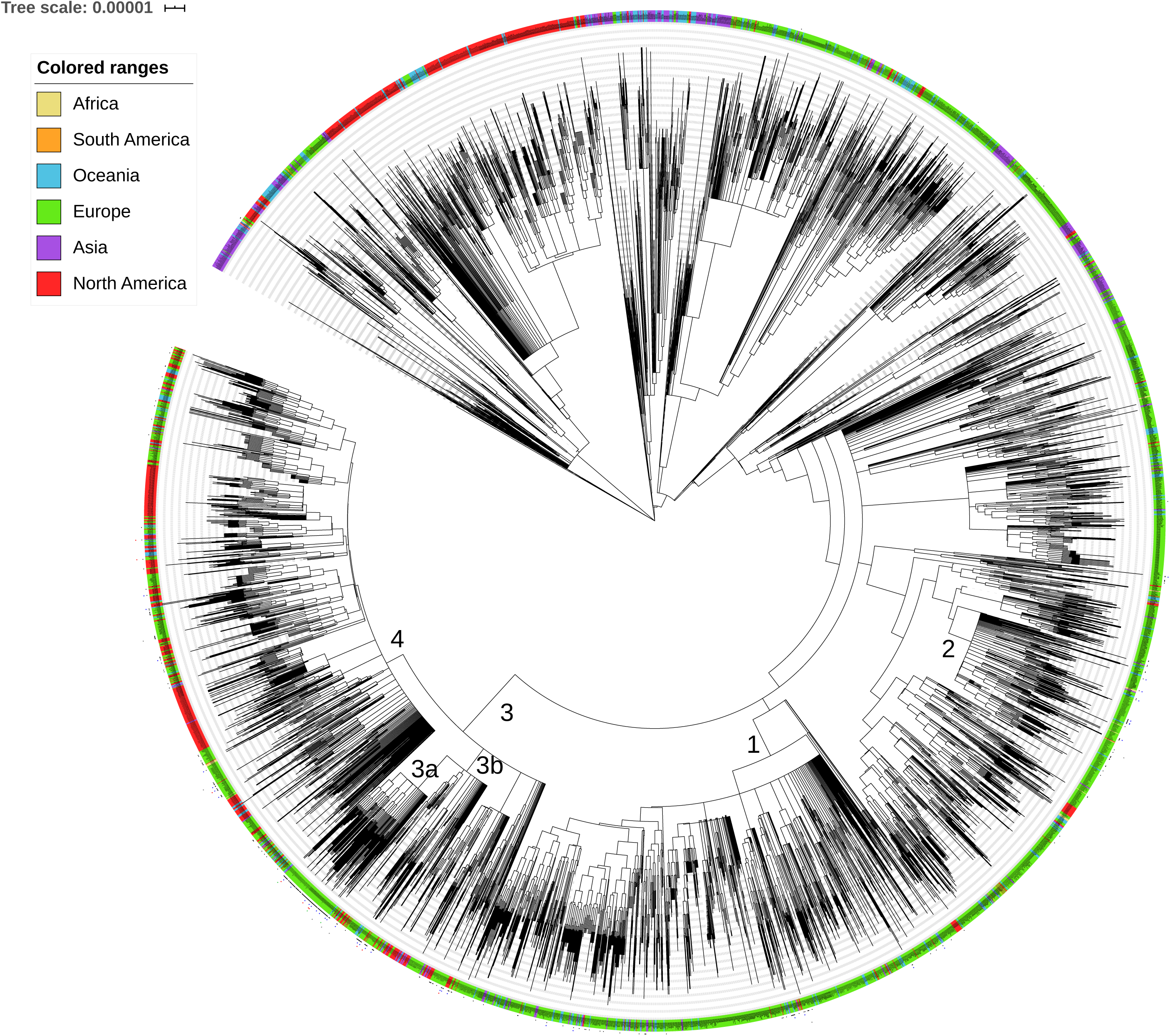
Phylogenetic tree of coronavirus worldwide. The phylogenetic tree was reconstructed from 5519 viral genomes collected around the world. Numbers (1 to 5) represent the cluster identified in the double hierarchical clustering. Red star corresponds to PClinO, blue star corresponds to PVirO, grey star corresponds to the loss of follow-up and green star corresponds to deceased patient.

Clinical outcomes of the 309 COVID-19-infected patients were investigated and integrated in our evolutionary analyses (Figure 1-3). Among these patients, we classified the outcomes in three categories including poor clinical outcome (PClinO, defined by either death or transfer to intensive care unit or hospitalization for 10 days or more), poor virological outcome (PVirO, defined by viral shedding persistence at day 10) and good clinical outcome (GO) (Supplementary Table 1). Coronavirus genomes isolates from 38 patients’isolates with PVirO were widely distributed across the groups, including diverse mutational events meaning that no correlation between higher viral loads and the genomic specificity of the virus was evidenced as previously reported (supplementary figure 2). For the 10 patients with PClinO, a majority of isolates were also distributed to all groups (supplementary figure 2). An exception concerned two patients (IHUCOVID-0318, IHUCOVID-0333) with one PClinO and one death that were clusterized together in a group of seven different isolates. A common nonsynonymous mutation in the membrane protein M (L129M, CTG>ATG, position 26907) was found in this cluster.

## DISCUSSION

Systematic gene level mutational analysis of the genomes allowed us to identify several unique features of the SARS-CoV-2 genome. In this study, we reported 315 substitution mutations including 126 synonymous mutations. Although these later mutations conserved AA, it has been reported that the fitness of RNA viruses with genomes harboring synonymous mutations could be affected^11^. Regarding the 178 nonsynonymous mutations, alteration due to the replacement of specific AA depends on several factors including their physicochemical properties, steric hindrance and the position of the AA in the tridimensional structure of the resulting protein. A key element of coronavirus host range is determined by the binding affinity between the spike S protein and the cellular receptor^12^. Here, thirty substitution mutations (including 18 nonsynonymous) were reported in the spike protein S. The spike protein mediates viral entry into host cells by recognizing angiotensin-converting enzyme 2 (ACE2) as its receptor^13^. We detected a missense mutation P330S (CCT>TCT) located at the border of the RBD^14^. Furthermore, a deletion of six nucleotides Δ21765-21770 removed two AA in the resulting protein. Mutations in the spike surface glycoprotein (A930V (24351C>T)) have already been reported in the Indian SARS-CoV-2^15^. All these mutations in protein S could influence host range and transmissibility of the virus. Consequently, the spike S glycoprotein is a potential target among potential therapeutic strategies. Focusing on the mutations identified in our study enables to drive the strategy to target special positions of the protein. In this line, specific antiviral therapies on S protein should be conducted *in vitro* (involving RBD-ACE2 blockers, S cleavage inhibitors, neutralizing antibodies, inhibitors and small interfering)^16^. Additionally, truncated proteins were also detected for ORF7a with a stop codon mutation (position 38 on 122 AA protein length) and for ORF3a with a frameshift mutation leading to an aborted protein missing 19 terminal AA. These mutations represent major changes in the protein architecture that have to be experimentally tested in order to assess the impact on protein function.

In evolutionary biology, convergent evolution is a central concept. The importance of a mutation is greatly strengthened if one can show that the fitness has evolved independently multiple times^17^. Two deletion events (Δ686-694; Δ1605-1607) were detected in three and two variants while the COVID-19-infected patients were not epidemiologically-related. These results strongly support a convergent evolution and could suggest convergent adaptation of the virus.

Phylogenetic trees were reconstructed by including isolates from France and all isolates worldwide. Isolates were grouped according to continents, but the large circulation of the virus was evidenced. Interestingly, we detected only two isolates belonging to the S-type in our 309 patients’ cohort^18^. Noteworthy, an almost exclusive group including SARS-CoV-2 genomes from 98 patients monitored in IHU Mediterranée Infection was phylogenetically close and regrouped in a clade of 105 genomes. Classification based on mutation profiles enables the inclusion of patients in specific groups and correlates this clustering with the clinical features in order to favor a better approach to treatments. We reported widespread variants that extent to different groups and pointed five clustering groups that spread to different lineages. Regarding the clinical evolution of patients, PClinO and PVirO were widely distributed in all groups identified in our clustering analysis as well as in the phylogenetic trees as previously observed^19^. A single exception was reported in a subgroup with a missense mutation in the gene encoding membrane protein M (L129M, CTG>ATG) in which two PClinO (including on death) were observed. This specific mutation in membrane protein M (L129M) has to be carefully investigated in future large cohort as potential determinant of clinical evolution. But at this stage, more data have to be integrated to state any conclusion or consequence about this mutation. Overall, these results suggested a clinical outcome that is not dependent on viral variants but rather a patient-dependent clinical outcome.

In conclusion, the mutational signatures and evolution of the coronavirus SARS-CoV-2 from 309 patients monitored in IHU Mediterranée Infection were unraveled. Inclusion of the French cohort allowed us to identify 161 novel mutations never reported before in coronavirus isolates worldwide. This mutations list represents a huge library of potential targets to test experimentally. The isolation and the genomic studies of the 309 SARS-CoV-2 genomes are paving the way for new diagnostic, therapeutic and prophylactic approaches.

## METHODS

### Patients

The study was conducted at Assistance Publique-Hôpitaux de Marseille (AP-HM), Southern France in the Institut Hospitalo-Universitaire (IHU) Méditerranée Infection (https://www.mediterranee-infection.com/). All data from patients were anonymized. The study was conducted on patients included from February 29^th^ to April 4^th^. Individuals with PCR-documented SARS-CoV-2 RNA from a nasopharyngeal sample^20^ were selected for genome sequencing. Viral loads ranged as follows: 93 patients with PCR Ct <16; 69 patients with [16<Ct<17]; 95 patients with [17<Ct<18]; 30 patients with [18<Ct<19]; 17 patients with [19<Ct<20] and 5 patients with Ct >20.

### Genome sequencing and assembling

From nasopharyngeal samples, genomic RNA was extracted using the EZ1 biorobot with the EZ1 DNA tissue kit (Qiagen, Hilden, Germany) as previously described ^21^. In the reverse transcription (RT) step, cDNA was reverse transcribed from total viral RNA samples using TaqMan Reverse Transcription (Life technologies applied biosystems) following the manufacturer’s recommendations. PCR program was set as follows: 25°C, 10 min, 48°C, 30 min, 95°C, 5 min, 4°C. DNA Polymerase I Large Klenow Fragment (BioLabs) was used for generating double stranded cDNA. 20 µL of each sample were added to a mix (10 µL) of: Buffer 2 Neb 10X, DNA polymerase large Klenow Fragment, dNTPs working solution (10mM) for 1 h at 37°C. cDNA were purified using beads Agencourt AMPure (Beckman Coulter) and then sequenced with the paired end strategy on a MiSeq sequencer (Illumina Inc, San Diego, CA,USA) with the Nextera XT DNA sample prep kit (Illumina). To prepare the paired-end library, the « tagmentation » step fragmented and tagged the DNA. Then, limited cycle PCR amplification (12 cycles) completed the tag adapters and introduced dual-index barcodes. After purification on AMPure XP beads (Beckman Coulter Inc, Fullerton, CA, USA), libraries were normalized on specific beads according to the Nextera XT protocol (Illumina). Normalized libraries were pooled into a single library for sequencing on MiSeq. The pooled single strand library was loaded onto the reagent cartridge and then onto the instrument along with the flow cell. Automated cluster generation and paired-end sequencing with dual index reads were performed in single 39-hours run in 2×250-bp as previously described ^21,22^. After sequencing, reads were mapped on the reference SARS-CoV-2 isolate Wuhan-Hu-1 (MN908947.3) using CLC Genomics workbench v.7 with the following thresholds: 0.8 for coverage and 0.9 for similarity. For each sample, detailed per-base coverage information was extracted and all mutations with a coverage ≥5 were considered.

### Hierarchical clustering

Data were ordered and grouped using the distance matrix computation and hierarchical clustering analysis calculated by MultiExperimentViewer^23^. A double clustering was carried out based on Pearson correlation as a distance matrix and complete linkage clustering as a linkage method.

### Sequence dataset

All genomes available on April 15 of SARS-CoV-2 were downloaded from GISAID (Global Initiative; https://www.gisaid.org/) with acknowledgment ^24,25^. All genomes were deposited at EMBL-EBI under the BioProject: PRJEB37722

AA: Amino Acid
GO: good clinical outcome
Indel: Insertion and deletion
ORF: Open Reading Frame
PClinO: poor clinical outcome
PVirO: poor virological outcome
RBD: Receptor Binding Domain
SD: Standard Deviation
SNP: Single Nucleotide Polymorphism
UTR: Untranslated Transcribed Region

## ETHICS APPROVAL AND CONSENT TO PARTICIPATE

The study was approved by the Ethical Committee of the University Hospital Institute Méditerranée Infection (N°: 2020-016). Informed consent was obtained from all patients. The study was performed in accordance to the good clinical practices recommended by the Declaration of Helsinki.

## AVAILABILITY OF DATA AND MATERIALS

All genomes were deposited at EMBL-EBI under the BioProject: PRJEB37722

## AUTHOR’S CONTRIBUTIONS

AL and DR supervised and conceived the project. AL designed the evolutionary studies and interpreted the data. AL and JD performed the experiments and analyzed the data. JCL, PEF, PC, AC, LB helped with data acquisition. AL wrote the manuscript. The authors read and approved the final manuscript.

## ACKNOWLEDGMENTS

Respectful thanks to all patients and their families enrolled in this study. The authors would like to thank all the engineers, technicians, and clinicians who contributed to this investigation. The authors gratefully acknowledge Olivia Ardizzoni, Vincent Bossi, Madeleine Carrara and Sofiane Regoui for help in sequencing the genomes. The authors acknowledge Yolande Obadia and Audrey Giraud-Gatineau for help in data acquisition. We gratefully thank the authors, the originating and submitting laboratories for their sequence and metadata shared through GISAID (Global Initiative; https://www.gisaid.org/). We acknowledge all the members of CoV-GLUE, Nextstrain (https://nextstrain.org/), and virological.org for sharing their analysis in real-time.

## FIGURES, TABLES AND ADDITIONAL FILES

**Supplementary Table 1. Baseline characteristics of the 309 COVID-19 infected patients**.

**supplementary figure 1. Emergence and phylogenetic tree of coronavirus in France**.

**supplementary figure 2. Phylogenetic tree of coronavirus from the cohort of 309 COVID-19 infected patients**. The phylogenetic tree was reconstructed from 309 viral genomes in this study. Numbers (1 to 5) represent the cluster identified in the double hierarchical clustering. The reference Wuhan coronavirus isolate was underlined in purple. Red star corresponds to PClinO, blue star corresponds to PVirO, grey star corresponds to the loss of follow-up and green star corresponds to deceased patient. Mutations were labeled along branches.

## AUTHOR INFORMATION

Correspondence to Didier Raoult or Anthony Levasseur *S: synonymous mutation; NS* : *nonsynonymous mutation*

