## Supplementary material for "Genomic diversity and evolution of coronavirus (SARS-CoV-2) in France from 309 COVID-19-infected patients"

**Supplementary Table 1. Baseline characteristics of the 309 COVID-19 infected patients.**

|  | **Total** | **Poor virological outcome** | **Poor clinical outcome** |
| --- | --- | --- | --- |
|  | **n (%)** | **n (%)** | **n (%)** |
| **Group size** | 309 (100%) | 38 (12.2%) | 10 (3.2%) |
| **Age (years)** |  |  |  |
| Mean (SD) | 49.6 (19.3) | 53.2(18.6) | 72 (16.28) |
| Median [Min-Max] | 48.0 [34-98] | 51.0 [42-89] | 75 [67-89] |

**Supplementary Figure 1. Emergence and phylogenetic tree of coronavirus in France.**

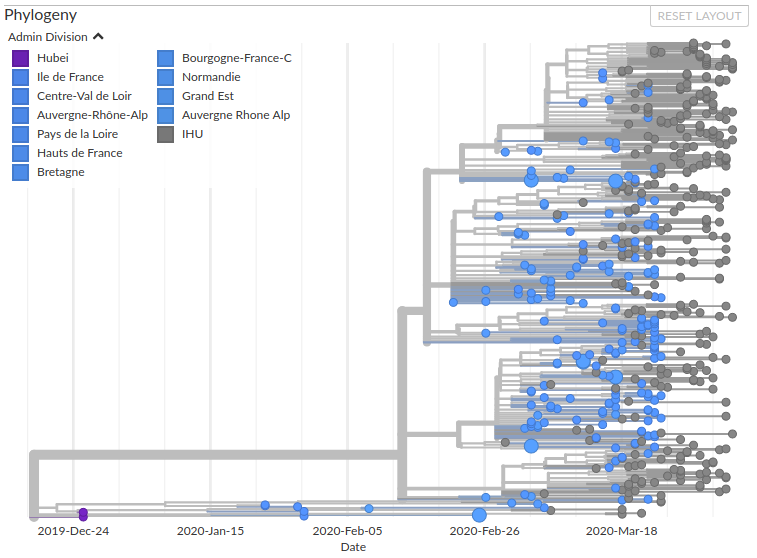

**Supplementary Figure 2. Phylogenetic tree of coronavirus from the cohort of 309 COVID-19 infected patients.** The phylogenetic tree was reconstructed from 309 viral genomes in this study. Numbers (1 to 5) represent the cluster identified in the double hierarchical clustering. The reference Wuhan coronavirus isolate was underlined in purple. Red star corresponds to PClinO, blue star corresponds to PVirO, grey star corresponds to the loss of follow-up and green star corresponds to deceased patient. Mutations were labeled along branches.

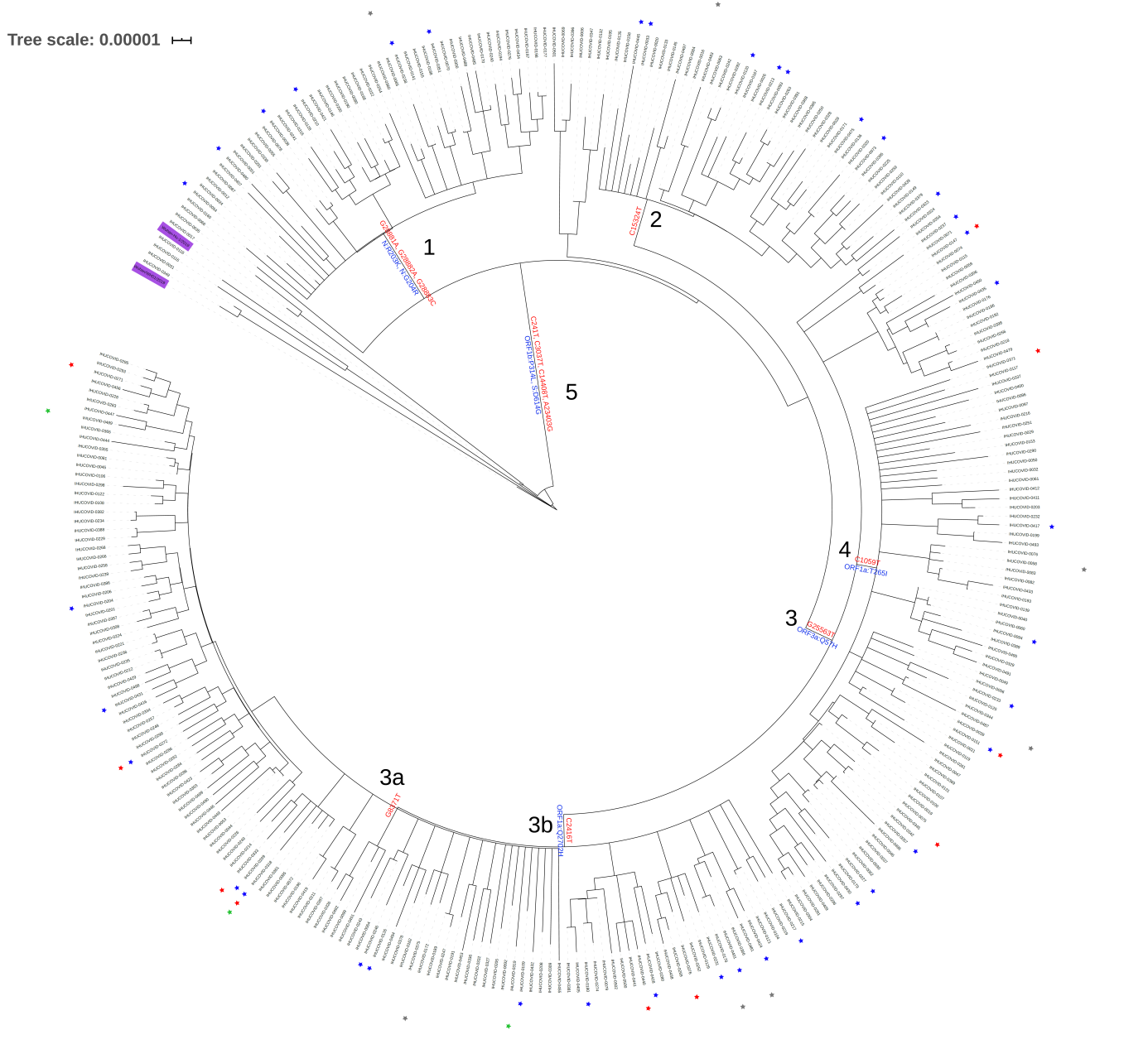
